# Elunetirom, a brain-targeted TRβ prodrug, promotes neuronal plasticity and mitochondrial biogenesis-related signaling in primary neuronal cultures

**DOI:** 10.64898/2026.07.21.739927

**Authors:** Jason R. Harris, Jill Baccei, Will Stratton, Stephen M. Stahl, Roger S. McIntyre, Thomas S. Scanlan, Gudarz Davar

## Abstract

Major depressive disorder and bipolar depression are disabling illnesses associated with impaired neuroplasticity, mitochondrial dysfunction, and altered cellular bioenergetics. Available pharmacotherapies often have delayed onset, incomplete efficacy, and limited effects on underlying plasticity and energetic systems. Non-selective thyroid hormone therapies provide evidence that enhancing central thyroid hormone signaling may improve depressive symptoms, but systemic cardiovascular, skeletal, and metabolic effects limit their broader use. Rapid-acting treatments such as esketamine and psychedelics further support neuroplasticity as a therapeutic strategy, although psychotomimetic, hallucinogenic, and implementation burdens may constrain widespread use. Elunetirom is a brain-targeted, fatty acid amide hydrolase-sensitive prodrug of LL-340001, a potent thyroid hormone receptor activator as demonstrated in transfected cell-line and brain-slice target-engagement assays. We evaluated elunetirom and LL-340001 in primary cortical and mature hippocampal neuronal cultures. LL-340001 increased MAP-2-positive neuron number, neurite length, branching, and neurite extremities in immature cortical neurons. In mature hippocampal neurons, elunetirom rapidly increased neurite network and synapse number within 24 hours, with effects persisting through 72 hours, while LL-340001 preserved synapse number following amyloid-β1–42 challenge. LL-340001 also increased nuclear PGC-1α and NRF2, the number of functional mitochondria measured by MitoTracker, and ATP content. Pharmacologic inhibition indicated that the LL-340001-induced increase in cortical MAP-2-positive neuron number was sensitive to TrkB inhibition but not 5-HT2A antagonism. Together, these findings indicate that elunetirom and LL- 340001 engage complementary neuroplasticity and mitochondrial biogenesis-related programs in neuronal systems and support further investigation of brain-targeted thyromimetic signaling as a mechanistically differentiated approach for major depressive disorder and bipolar depression.

## Introduction

Major depressive disorder (MDD) and bipolar depression remain leading causes of disability, morbidity, and recurrent functional decline, and are associated with substantial symptom heterogeneity, chronicity, and functional impairment that complicate personalized management and limit recovery for many patients [1–4]. Conventional treatments for mood disorders have suboptimal overall efficacy, often provide inadequate improvement in disabling symptom domains such as anhedonia, have delayed time-to-peak efficacy, and are associated with relatively low rates of remission and sustained psychosocial recovery [1–4]. Moreover, the best-established augmentation approaches in MDD include atypical antipsychotics, which are associated with agent-specific tolerability and safety concerns [5].

During the past two decades, a confluence of data has supported a disease model of mood disorders that implicates alterations in bioenergetics, neuroplasticity, and impaired neural circuits/network activity [6–9]. According to this mechanistic model, depressive syndromes are a result of impaired structure and function in discrete corticolimbic networks and cellular energetic dysfunction in networks that fail to maintain synaptic connectivity, adaptive plasticity, and metabolic resilience under chronic stress and recurrent illness burden.

Amongst the circuits and nodal regions, the prefrontal cortex has emerged as one of the brain regions most consistently impaired in MDD, with structural and functional abnormalities linked to cognitive, motivational, and affective dysfunction [6]. In parallel, reduced hippocampal gray matter volume has been identified across major depression and bipolar disorder, supporting the concept that depressive illness is associated with measurable abnormality of the circuits subserving stress adaptation, emotional regulation, and memory integration [7]. Although the directionality of these associations is complex, the cumulative evidence strongly supports the view that MDD and bipolar depression are characterized by network-level atrophy or impaired maintenance of neuronal architecture within prefrontal-hippocampal systems.

At the cellular level, mitochondrial and bioenergetic dysfunction are increasingly implicated in MDD and BD [8, 9]. Bioenergetic alterations have been observed across psychiatric disorders, but in mood disorders, bioenergetic and plasticity alterations are highly reproducible preclinical and clinical observations. In MDD, abnormalities in oxidative phosphorylation, ATP generation, oxidative stress handling, and mitochondrial metabolism have been implicated in illness biology [8]. These findings support the development of therapeutics centered on reversing impaired plasticity, stress vulnerability, and reduced circuit resilience.

Studies related to bipolar disorder have also identified alterations in energy production, redox imbalance, calcium dysregulation, and broader mitochondrial instability [9]. These observations are important because neurite outgrowth, dendritic remodeling, synaptogenesis, and sustained neurotransmission are energetically demanding processes that depend on intact mitochondrial trafficking and bioenergetic reserve, with PGC-1α serving as a master regulator of mitochondrial biogenesis and a nodal regulator of energy expenditure [10, 11]. A derivative of the robust evidence base as it pertains to bioenergetics and plasticity is the viable and testable hypothesis that restoring mitochondrial function while simultaneously driving structural plasticity would therefore mechanistically target two highly convergent pathophysiologic processes in MDD and bipolar depression.

Thyroid hormone signaling has long been linked to mood regulation, and non-selective thyroid hormone therapies have repeatedly shown therapeutic utility in MDD and bipolar disorder [12, 13]. In MDD, liothyronine (T3) has been shown to accelerate antidepressant response and augment treatment in refractory depression. In bipolar disorder, adjunctive levothyroxine (T4), often at supraphysiologic doses, has shown benefit in refractory bipolar depression and rapid-cycling illness. Importantly, these clinical signals are mechanistically plausible: thyroid hormones exert major effects on neuronal differentiation, maturation, synaptogenesis, and myelination, while also regulating mitochondrial activity, oxygen consumption, and mitochondrial biogenesis [14–19]. Consequently, thyroid action in the brain is simultaneously integral to two biological deficits: impaired neuroplasticity and impaired bioenergetics.

Despite this promise, thyroid hormone treatment remains infrequently prescribed in routine psychiatric practice [12, 13]. Acceptability of non-selective thyroid treatment in persons with MDD and bipolar depression is hampered by tolerability and safety concerns. For example, even where tolerability has been deemed acceptable in specialist settings, the requirement for monitoring and the perceived risk of cardiovascular, skeletal, neuropsychiatric, and metabolic adverse effects continue to limit widespread use of thyroid hormone as augmentation therapy in either MDD or bipolar depression by psychiatrists and other physicians.

Recent treatment discovery and development efforts in depression have increasingly prioritized interventions capable of producing rapid neuroplastic and neurotrophic effects, thereby enhancing synaptic organization and accelerating symptomatic improvement. Esketamine provides an approved proof-of-concept for this strategy, and investigational agents such as psilocybin further support the view that rapidly acting pro-plastic interventions may offer meaningful clinical benefit [20, 21]. Oral dextromethorphan-bupropion further illustrates the growing interest in rapid-acting, non-monoaminergic antidepressants, and mechanistic discussions have linked its antidepressant effects to NMDA receptor modulation, sigma-1 receptor signaling, and related downstream biology relevant to plasticity and trophic signaling [22–24].

However, dextromethorphan-bupropion is not approved for bipolar depression, and prescribing information highlights important limitations including activation of mania or hypomania, seizure risk, increased blood pressure, and neuropsychiatric adverse reactions [25]. More broadly, several rapidly acting agents remain concentrated in more difficult-to-treat or later-stage depressive illness and may still carry treatment-limiting safety, tolerability, or implementation burdens [20, 26, 27]. Accordingly, there remains a need for scalable, mechanistically informed pro-plastic and pro-trophic treatments that can be deployed across a broader range of patients, including those outside treatment-resistant populations, while avoiding treatment-limiting safety concerns and barriers to widespread use.

These converging lines of evidence support a clear therapeutic hypothesis: a thyromimetic delivered selectively to the CNS could preserve the mitochondrial, synaptogenic and clinical benefits of thyroid hormone for depression treatment while minimizing the peripheral liabilities that have limited adoption of T3 and T4 [28–30]. Prior work with CNS-directed thyromimetic strategies has already shown that selective thyroid hormone receptor agonism and brain-targeted prodrug approaches can enhance CNS thyroid hormone action while reducing peripheral exposure, thereby providing a translational precedent for brain-focused thyroid hormone pharmacology. For mood disorders characterized by impaired neural plasticity and bioenergetic insufficiency, such a strategy may represent a mechanistically differentiated approach to treatment beyond incremental symptomatic control [6–9].

The present study investigated a potent thyromimetic compound in primary neuronal culture systems designed to capture complementary dimensions of neuronal remodeling and energetic recovery. Specifically, we test the hypothesis that this compound engages mitochondrial biogenesis-related signaling through upregulation of PGC-1α and promotes neuroplastic change through BDNF-TrkB-linked signaling [10, 11]. We aimed to demonstrate increased MAP-2-positive neuron number and neuritogenesis in primary cortical neurons, increased neuritogenesis and synaptogenesis in mature hippocampal neurons, and improved mitochondrial abundance and function, as reflected by an increased number of functional mitochondria measured by MitoTracker, increased ATP content, and increased PGC-1α and NRF2 protein levels. In doing so, we sought to establish a mechanistic foundation for CNS-selective thyromimetic therapy as a mechanistically differentiated therapeutic approach for MDD and bipolar depression (Figure 1).

**Figure 1.**
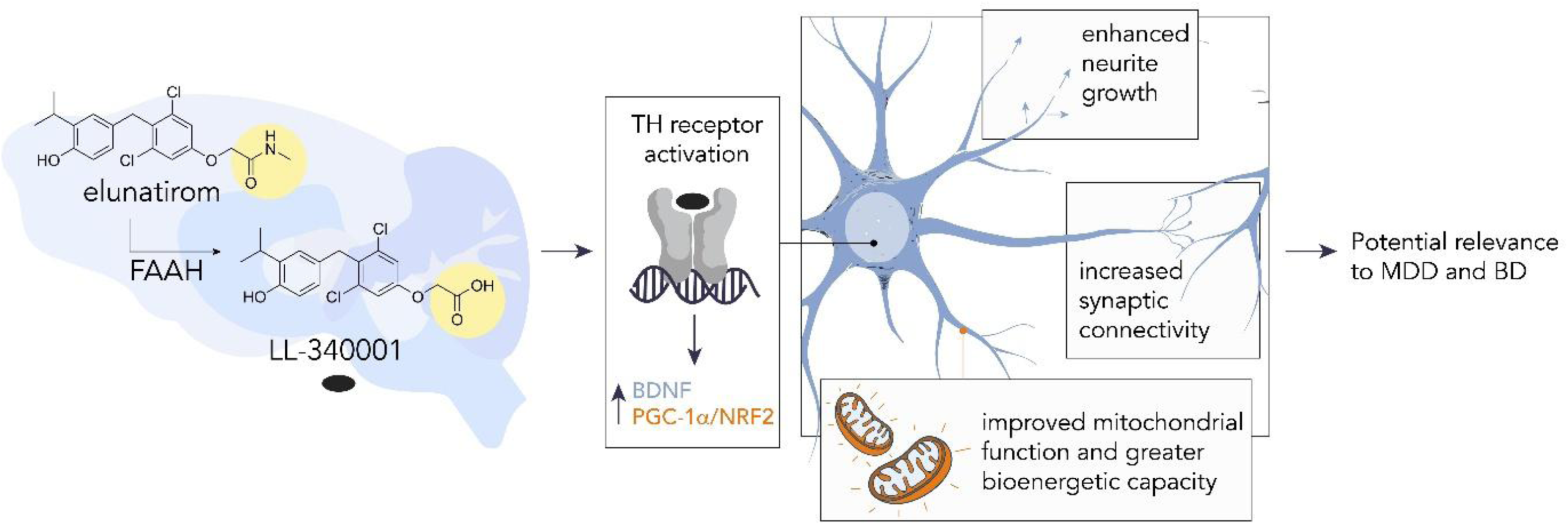
Proposed integrated mechanism of elunetirom and LL-340001 in primary neuronal cultures. Elunetirom is converted by fatty acid amide hydrolase (FAAH) to the active thyromimetic LL-340001, which activates thyroid hormone receptors and engages BDNF- and PGC-1α/NRF2-associated signaling. These pathways promote neurite growth, synaptic connectivity, and mitochondrial bioenergetic function, supporting potential relevance to major depressive disorder and bipolar depression.

## Materials and Methods

### TRα and TRβ reporter assays

Human TRα and TRβ reporter assays were performed in HEK293-derived cells expressing Gal4 DNA-binding domain/receptor fusion constructs together with a Gal4-responsive firefly luciferase reporter. Cells were equilibrated for 4 h in assay medium containing charcoal-stripped serum, then pretreated with vehicle (0.1% DMSO) or the selective FAAH inhibitor PF-3845 (1 μM) [31] for 30 min before addition of test compounds or triiodothyronine (T3). Compounds were incubated for 23 h at 37°C in 5% CO₂ at a final DMSO concentration of 0.15%. Luciferase activity was then measured by plate luminometry. T3 served as the reference agonist, and responses were normalized to the maximal T3 signal within each assay. Concentration-response curves were fit by nonlinear regression using GraphPad Prism, and EC50-equivalent values were interpolated at 50% of the normalized T3 response.

### FAAH hydrolysis

Elunetirom (ABX-002) was incubated with recombinant human FAAH, and formation of ABX-002A (LL-340001) was measured in the presence or absence of PF-3845 (1 μM). Reactions were performed in Tris-EDTA buffer (pH 8.0) containing bovine serum albumin. At 0–60 min, aliquots were quenched with methanol/acetonitrile containing tolbutamide, centrifuged, and supernatants were analyzed by LC-MS/MS. ABX-002A formation was quantified as peak area ratio relative to internal standard, and formation rates were determined from the slope of peak area ratio versus time.

### Organotypic mouse brain-slice cultures

Organotypic forebrain slice cultures were prepared from postnatal day 9 C57BL/6 mice born to timed-pregnant dams obtained from Charles River Laboratories. Pups were not selected by sex, and the brain-slice cultures were considered mixed sex. Animal procedures were conducted at Explora BioLabs under a protocol approved by the Explora BioLabs Institutional Animal Care and Use Committee (IACUC). Coronal slices (∼400 μm) were placed on culture inserts, maintained at 37°C in 5% CO₂, and allowed to recover for 72 h before treatment. Elunetirom and ABX-002A were added at the indicated concentrations (0.1% final DMSO) for 120 h, with media and compound replenished every 48 h. Slices were then collected for RNA isolation, and gene expression was quantified using a custom NanoString PlexSet panel. Raw counts were normalized in nSolver using an internal calibrator sample and the geometric mean of housekeeping genes (Gusb, Ppib, and Txn2). Fold changes relative to DMSO-treated controls were determined for the thyroid hormone-responsive genes iodothyronine deiodinase 3 (Dio3), hairless (Hr), Krüppel-like factor 9 (Klf9), and secreted frizzled-related protein 5 (Sfrp5); log2-transformed values were averaged to generate a composite response score. Concentration-response curves were fit in GraphPad Prism to derive EC50 values.

### Primary culture of cortical neurons

Primary cortical cultures were prepared from pooled cortices collected from embryonic day 15 (E15) Wistar rat fetuses as described by Callizot et al. [32]. Briefly, cortices at embryonic day 15 were dissected in L15 Leibovitz medium (Dutscher) containing 2% penicillin (10,000 U/mL) / streptomycin (10 mg/mL) (PS) solution (Dutscher) and 1% bovine serum albumin (BSA, Dutscher). Cortices were dissociated by trypsinization (0.05% trypsin and 0.02% ethylenediamine tetra-acetic acid (EDTA), Dutscher), and the reaction was stopped by the addition of Dulbecco’s modified Eagle medium (DMEM, Dutscher) containing grade II DNAse I (0.2 mg/mL, Dutscher) and 10 % fetal bovine serum (FBS, Fisher Scientific). Cells were mechanically dissociated and resuspended in Neurobasal™ medium (Fisher Scientific) supplemented with 2 % B27 (Fisher Scientific) containing 2 mM of L-glutamine (Fisher Scientific), 2% PS solution, and 10 ng/mL of brain-derived neurotrophic factor (BDNF, Tebubio). Medium was replaced every 48 hours.

### Primary culture of hippocampal neurons

Primary hippocampal cultures were prepared from pooled hippocampi collected from embryonic day 17 (E17) Wistar rat fetuses as described by Callizot et al. [33]. Briefly, hippocampi at embryonic day 17 were dissected in L15 Leibovitz medium (Dutscher) containing 2 % penicillin (10,000 U/mL) / streptomycin (10 mg/mL) (PS) solution (Dutscher) and 1% BSA (Dutscher). Hippocampi were dissociated by trypsinization (0.05% trypsin and 0.02% EDTA, Dutscher), and the reaction was stopped by the addition of DMEM with 4.5 g/L of glucose (Dutscher), containing grade II DNAse I (0.2 mg/mL, Dutscher), and 10% FBS (Fisher Scientific). Cells were mechanically dissociated and resuspended in Neurobasal™ medium (Fisher Scientific) supplemented with 2% B27 (Fisher Scientific) containing 2 mM of L-glutamine (Fisher Scientific), 2% PS solution, and 10 ng/mL of BDNF (Tebubio). Medium was replaced every 48 hours.

### Animal welfare and sex as a biological variable

For the organotypic mouse brain-slice study, timed-pregnant C57BL/6 dams were obtained from Charles River Laboratories. Pups were not selected by sex, and the brain-slice cultures were considered mixed sex. Animal procedures were conducted at Explora BioLabs under a protocol approved by the Explora BioLabs IACUC. For the primary neuronal culture studies, pregnant Wistar rats were housed and handled at Neuro-Sys under veterinary authorization C1301310. Fetal sex was not determined, and tissue from multiple fetuses was pooled before plating; accordingly, the cortical and hippocampal cultures were considered mixed-sex cultures. Because the Neuro-Sys studies involved in vitro cultures derived from embryonic tissue and did not constitute animal procedures requiring project authorization under the applicable European regulatory framework, separate institutional animal ethics approval was not required for the culture experiments. All procedures involving animals were conducted in accordance with the National Research Council (US) Guide for the Care and Use of Laboratory Animals and applicable European Union regulations, including Directive 2010/63/EU. The ARRIVE guidelines were considered not applicable to the in vitro embryonic neuronal culture experiments because they did not include in vivo animal procedures.

### In vitro LL-340001 treatments

DIV1-3 cortical neurons were incubated with LL-340001 (1 nM - 3 μM) or 0.1% dimethyl sulfoxide (DMSO, Dutscher) for 72 hours. Medium containing treatment compounds was replaced after 48 hours. BDNF (50 ng/mL in PBS, Dutscher) was applied as a positive control.

### Mechanistic antagonist treatments in immature cortical neurons

To evaluate the contribution of TrkB and 5-HT2A signaling to LL-340001-induced cortical plasticity, DIV1 primary rat cortical cultures were seeded at 25,000 cells per well in poly-L-lysine- and collagen-coated 96-well plates and pretreated for 1 h with the TrkB antagonist ANA-12 (100 nM), the 5-HT2A antagonist M100907 (50 nM), or vehicle. LL-340001 (3 or 30 nM), psilocin (300 nM), BDNF (50 ng/mL), or vehicle (0.1% DMSO) was then added, and cultures were maintained for 72 h. All vehicle and test-compound conditions were maintained in the standard culture medium containing 10 ng/mL BDNF; the 50 ng/mL BDNF condition served as an established positive control selected to produce an optimal increase in MAP-2-positive neuron number. Antagonist-only and test-compound-plus-antagonist conditions were included. Six wells were assigned per condition. After treatment, cells were fixed in 95% ethanol/5% acetic acid for 15 min at -20°C and immunostained with anti-MAP-2 antibody (1:400). Twenty-five fields per well, representative of the whole-well area, were acquired at 20× magnification using an Operetta CLS high-content imaging system and analyzed using Harmony software to quantify MAP-2-positive neuron number, total neurite length, neurite roots, and neurite extremities. Wells affected by documented technical failures, including absence of cells or inadequate staining, were excluded before analysis, resulting in n = 5-6 wells per condition.

### In vitro elunetirom and LL-340001 treatments

DIV17-18 or DIV17-20 hippocampal neurons were incubated with elunetirom (0.3 nM – 1 μM), LL-340001 (1 nM - 3 μM), or 0.1% DMSO (Dutscher) for 24 or 72 hours. Medium containing treatment compounds was replaced after 48 hours. BDNF (50 ng/mL (Tebubio) in PBS (Dutscher)) or psilocin (300 nM (Sigma-Aldrich) in acetonitrile (Dutscher)) was applied as a positive control.

### In vitro Aβ1–42 and LL-340001 treatment

DIV17-18 hippocampal neurons were incubated with LL-340001 (1 nM - 1 μM) or 0.1% DMSO (Dutscher) for 4 hours before application of 1.25 μM Aβ1-42 (4147158, Bachem; prepared as described by Callizot et al. [33]), for 24 hours. BDNF (50 ng/mL in PBS, Dutscher) in acetonitrile (Dutscher)) was applied as a positive control.

### MitoTracker labeling

DIV20 supernatants were removed, and viable hippocampal cells were incubated with 500 nM Mitotracker™ Deep Red FM (M22426, Thermofisher) in 1x Hank’s Balanced Salt Solution (HBSS, cod. 15266355, Fisher Scientific). Hippocampal cells were then fixed and immunostained (see section 2.12.)

### Immunostaining and image analysis

The cells were fixed in 95% ethanol/5% acetic acid (Sigma-Aldrich), washed with PBS, and blocked and permeabilized in PBS containing 1% saponin (Sigma-Aldrich) and 1% FBS (Fisher Scientific). Primary antibodies were MAP-2 (1:1000; catalog no. ab5392, Abcam), synaptophysin (SYN; 1:500; catalog no. 101008, Synaptic Systems), PSD95 (1:50; catalog no. 124 011, Synaptic Systems), PGC-1α (1:100; catalog no. NB100-60955, Novus Biologicals), and NRF2 (1:500; catalog no. PA5-27882, Thermo Fisher Scientific). Secondary antibodies were anti-mouse (1:400; catalog nos. SAB400042, Sigma-Aldrich, and 715-545-151, Jackson ImmunoResearch), anti-chicken (1:400; catalog nos. SAB4600179 or SAB4600039, Sigma-Aldrich, and 703-605-155, Jackson ImmunoResearch), anti-rabbit (1:400; catalog nos. SAB4600084, Sigma-Aldrich, and 711-605-152, Jackson ImmunoResearch), and anti-goat (1:400; catalog no. 705-575-147, Jackson ImmunoResearch). Nuclei were counterstained with Hoechst 33258 (1:1000; Sigma-Aldrich). Immunofluorescent images were acquired using an Operetta CLS high-content imaging system (Revvity) at 20× or 40× magnification and analyzed using Harmony software (Revvity). Synapse number per neuron was operationally defined as the number of colocalized SYN- and PSD95-positive puncta associated with MAP-2-positive neuronal structures, normalized to the number of MAP-2-positive neurons. This immunocytochemical endpoint reflects structurally identified synaptic contacts rather than electrophysiologic function. For each well, measurements from all acquired image fields were aggregated to generate a single well-level value; individual image fields and cells were treated as subsamples and were not analyzed as independent experimental units.

### ATP content measurement

DIV20 supernatants were removed, and ATP content was assessed in viable hippocampal cultures using an ADP/ATP Ratio Assay Kit (MAK135, Sigma-Aldrich). Bioluminescent signal was detected using GlowMax® Discover (Promega).

### Statistics

Data are presented as mean ± standard error of the mean (SEM). For the Neuro-Sys cortical and hippocampal neuronal culture assays, the individual culture well was the experimental unit. Measurements from all image fields acquired within a well were aggregated to produce one value per well; image fields and individual cells were considered subsamples. Because cells from multiple fetuses were pooled before plating, the reported n values denote replicate wells per condition from pooled mixed-sex cultures rather than individual animals. For the mouse brain-slice gene-expression assay, n denotes individually analyzed brain-slice samples per treatment concentration. For the TRα and TRβ reporter assays, n denotes assay determinations. Statistical analysis was performed using GraphPad Prism 111 (GraphPad Software Inc., USA). A one-way ANOVA followed by Fisher’s LSD test for multiple comparisons was applied for all assays. Statistical significance was assumed at p < 0.05.

## Results

### Elunetirom is a prodrug for the receptor-active thyromimetic LL-340001 and becomes functionally equivalent to LL-340001 in neural tissue

We first sought to define the pharmacologic relationship between elunetirom and its active metabolite, the thyroid hormone receptor agonist LL-340001. In direct thyroid hormone receptor reporter assays, LL-340001 displayed potent agonist activity at both TRα and TRβ, whereas elunetirom itself was weak, exhibiting less than one-tenth the potency of LL-340001 and only partial efficacy relative to the reference agonist T3 (Fig. 2). These findings established LL-340001 as the receptor-active species and suggested that any robust activity of elunetirom in neuronal systems would likely require metabolic conversion to the active metabolite.

**Fig. 2.**
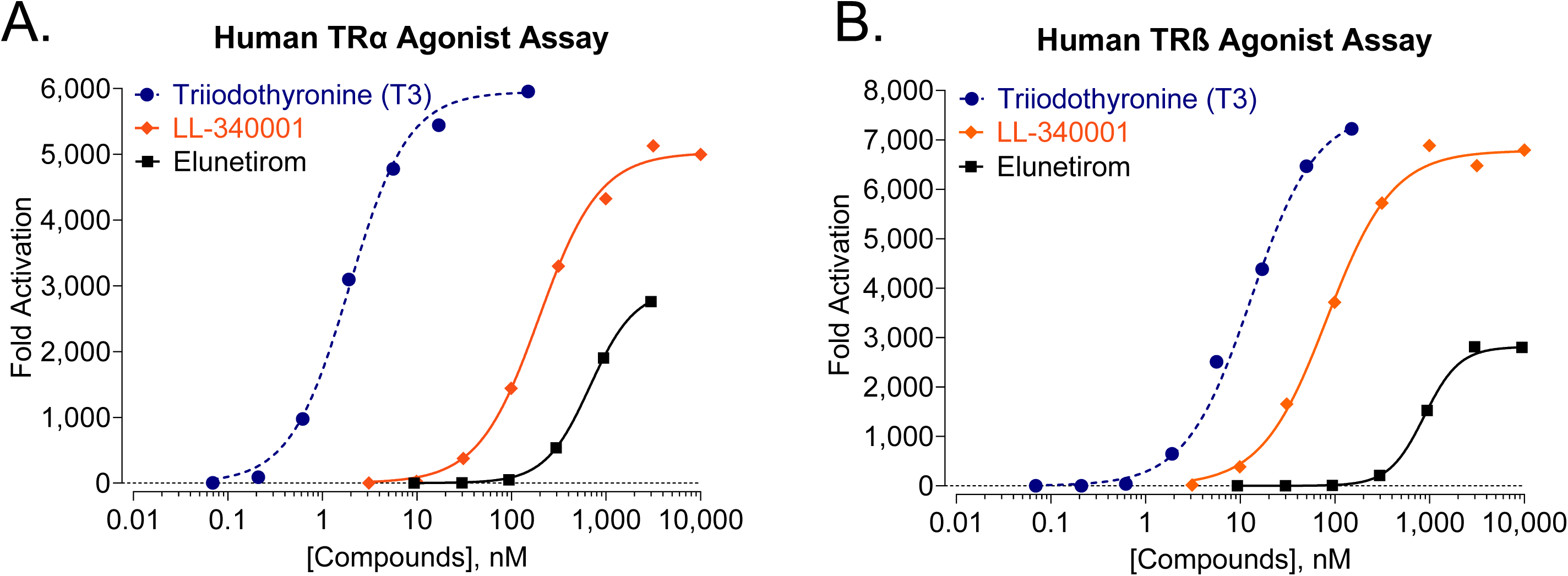
Agonist activity of elunetirom and LL-340001 at human TRα and TRβ relative to triiodothyronine. Mean (n=2) agonist activity of elunetirom, LL-340001, and T3 against human TRα (A) and TRβ (B). Elunetirom TRα EC_50_ = 677 nM (max activation 44%); elunetirom TRβ EC_50_ = 876 nM (max activation 39%); LL-340001 TRα EC_50_ = 196 nM (max activation 94%); LL-340001 TRβ EC_50_ = 83 nM (max activation 80%); T3 TRα EC_50_ = 1.9 nM (max activation 100%); T3 TRβ EC_50_ = 13 nM (max activation 100%). EC50-equivalent values were interpolated at 50% of the normalized T3 response. Elunetirom TRα T3-50 = >30,000 nM (max activation 44%); elunetirom TRb T3-50 = >30,000 nM (max activation 39%); LL-340001 TRα T3-50 = 280 nM (max activation 80%); LL-340001 TRβ T3-50 = 100 nM (max activation 94%).

**Table 1.** Rates of conversion of elunetirom.

| Elunetirom | Slope based on the formation of LL-340001 |  |
| --- | --- | --- |
|  | hFAAH |  |
| + No inhibitor | 0.00169 |  |
| + 1 μM PF-3845 | 0.00 | |
| inhibition | 100% |  |

**Fig. 3.**
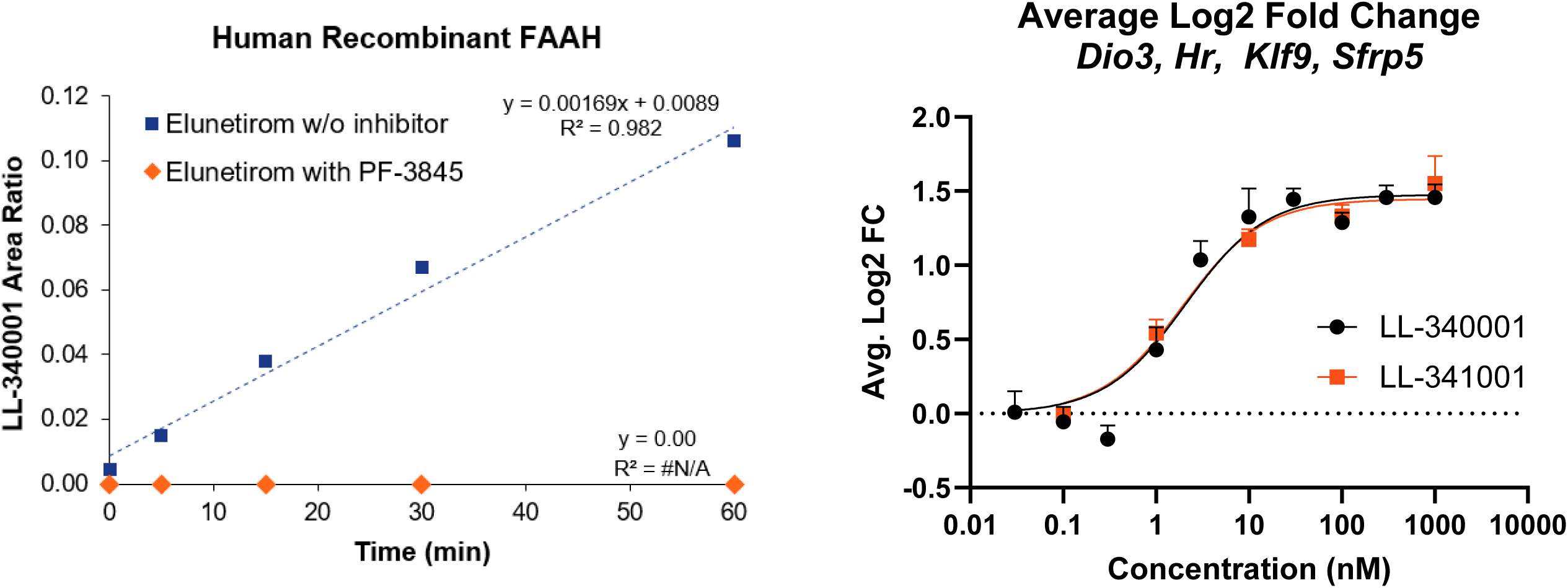
FAAH-dependent conversion of elunetirom to LL-340001 and functional equivalence in mouse brain slices. Effects of elunetirom and LL-340001 on gene expression in cortical brain slices from perinatal mice in culture. AveLog2FC EC50 values were 1.93 and 2.08 nM for elunetirom and LL-340001, respectively. Data points are mean ± SEM. N=5/treatment concentration; *Dio3*=deiodinase 3; *Hr*=hairless; *Klf9=* Krueppel-like factor; SEM=standard error of the mean; *Sfrp5*=secreted frizzled related protein 5.

**Figure 4.**
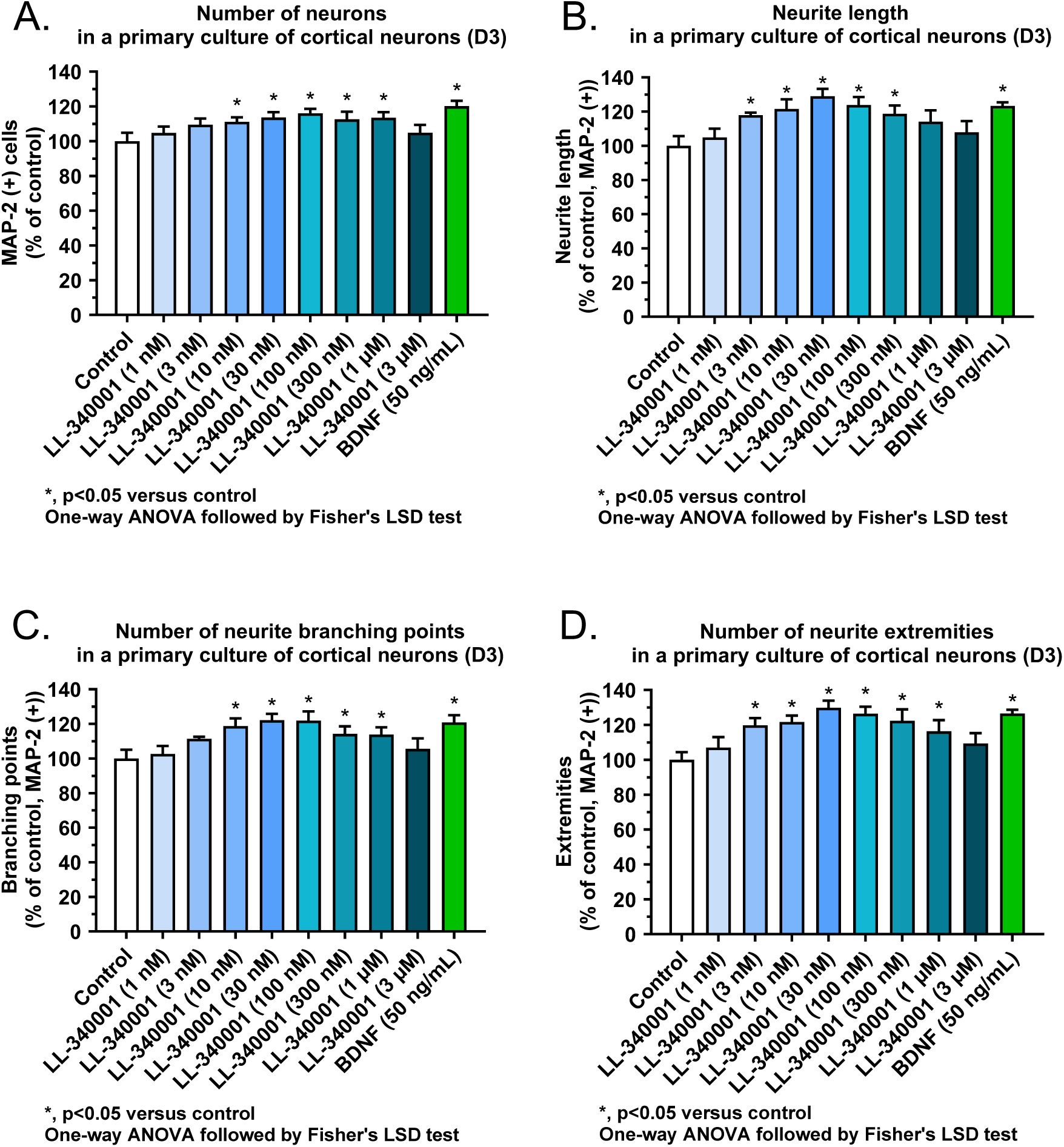
LL-340001 increases MAP-2-positive neuron number and neuritic complexity in immature primary cortical neurons. Effect of LL-340001 on the number of MAP-2 (+) neurons (A), neurite network (B), branching points (C), and neurite extremities (D) in a primary culture of cortical neurons (day 3). LL-340001 was assessed at 1 nM, 3 nM, 10 nM, 30 nM, 100 nM, 300 nM, 1 μM, and 3 µM, as well as the positive control, BDNF (50 ng/mL). Results are expressed as a percentage of control condition as mean ± SEM (n = 5-6). Data were compared using a one-way ANOVA followed by Fisher’s LSD test. *p< 0.05 versus control was considered significant.

**Figure 5:**
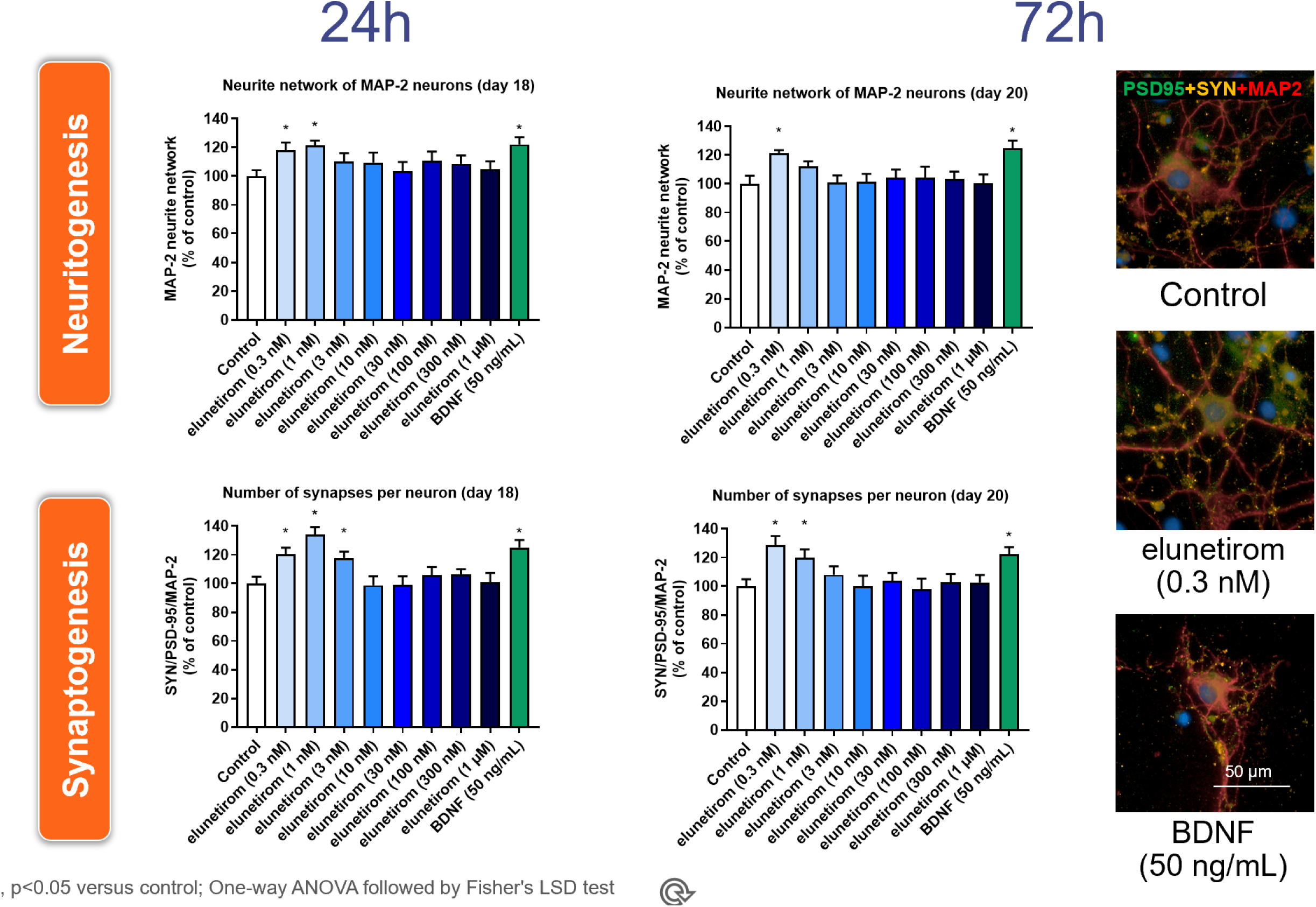
Elunetirom rapidly increases neurite network and synapse number in mature primary hippocampal neurons, with effects sustained through 72 hours. Effect of a 24-hour or 72-hour treatment with Elunetirom on the number of MAP-2 (+) neurons (A); their neurite network (B), and the number of synapses (C) in a primary culture of hippocampal neurons (day 18 or day 20). Elunetirom was assessed at 0.3 nM, 1 nM, 3 nM, 10 nM, 30 nM, 100 nM, 300 nM, and 1 μM, along with the positive control BDNF (50 ng/mL). Results are expressed as a percentage of control condition as mean ± SEM (n = 4-6). A one-way ANOVA followed by Fisher’s LSD test was used to compare all the experimental conditions. *p< 0.05 versus control was considered significant.

**Fig. 6.**
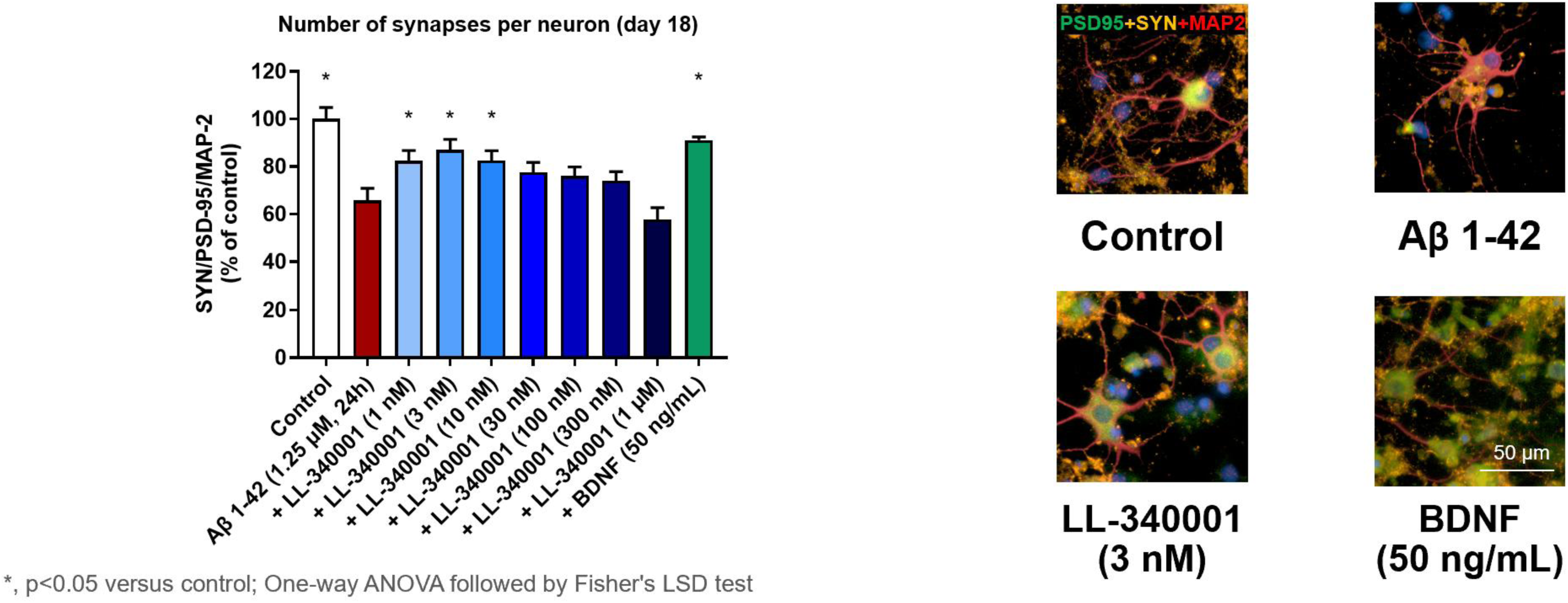
LL-340001 preserves synapse number in Aβ1-42–challenged mature primary hippocampal neurons. Effect of LL-340001 on the number of MAP-2 (+) the number of synapses in a primary culture of hippocampal neurons (day 18) injured with Aβ1-42 (1.25 μM, 24 hours). LL-340001 was assessed at 1 nM, 3 nM, 10 nM, 30 nM, 100 nM, 300 nM and 1 μM, as well as the positive control, BDNF (50 ng/mL). Results are expressed as a percentage of control condition as mean ± SEM (n = 5-6). A one-way ANOVA followed by Fisher’s LSD test was used to compare all the experimental conditions. *p< 0.05 versus Aβ1-42 was considered significant.

**Fig. 7.**
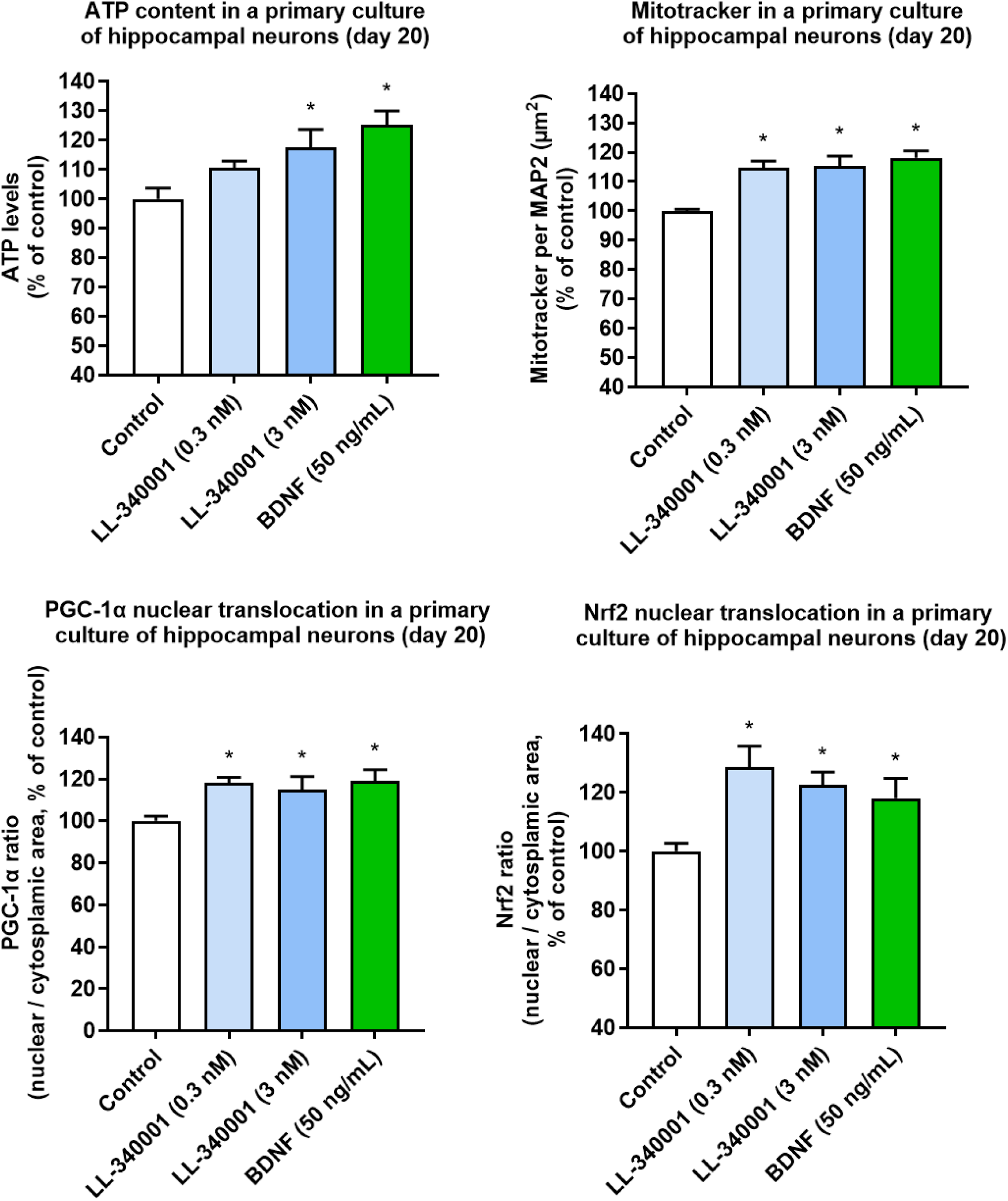
LL-340001 enhances mitochondrial biogenesis-associated signaling and bioenergetic output in mature primary hippocampal neurons. Effect of LL-340001 on ATP content (A), the number of functional mitochondria (B), and nuclear translocation of transcription factors PGC-1α (C) and Nfr2 (D) in a primary culture of hippocampal neurons (day 20). LL-340001 was assessed at 0.3 nM and 3 nM along with positive control, BDNF (50 ng/mL). Results are expressed as a percentage of control condition as mean ± SEM (n = 4-6). A one-way ANOVA followed by Fisher’s LSD test was used to compare all the experimental conditions. *p< 0.05 versus control was considered significant.

**Fig. 8.**
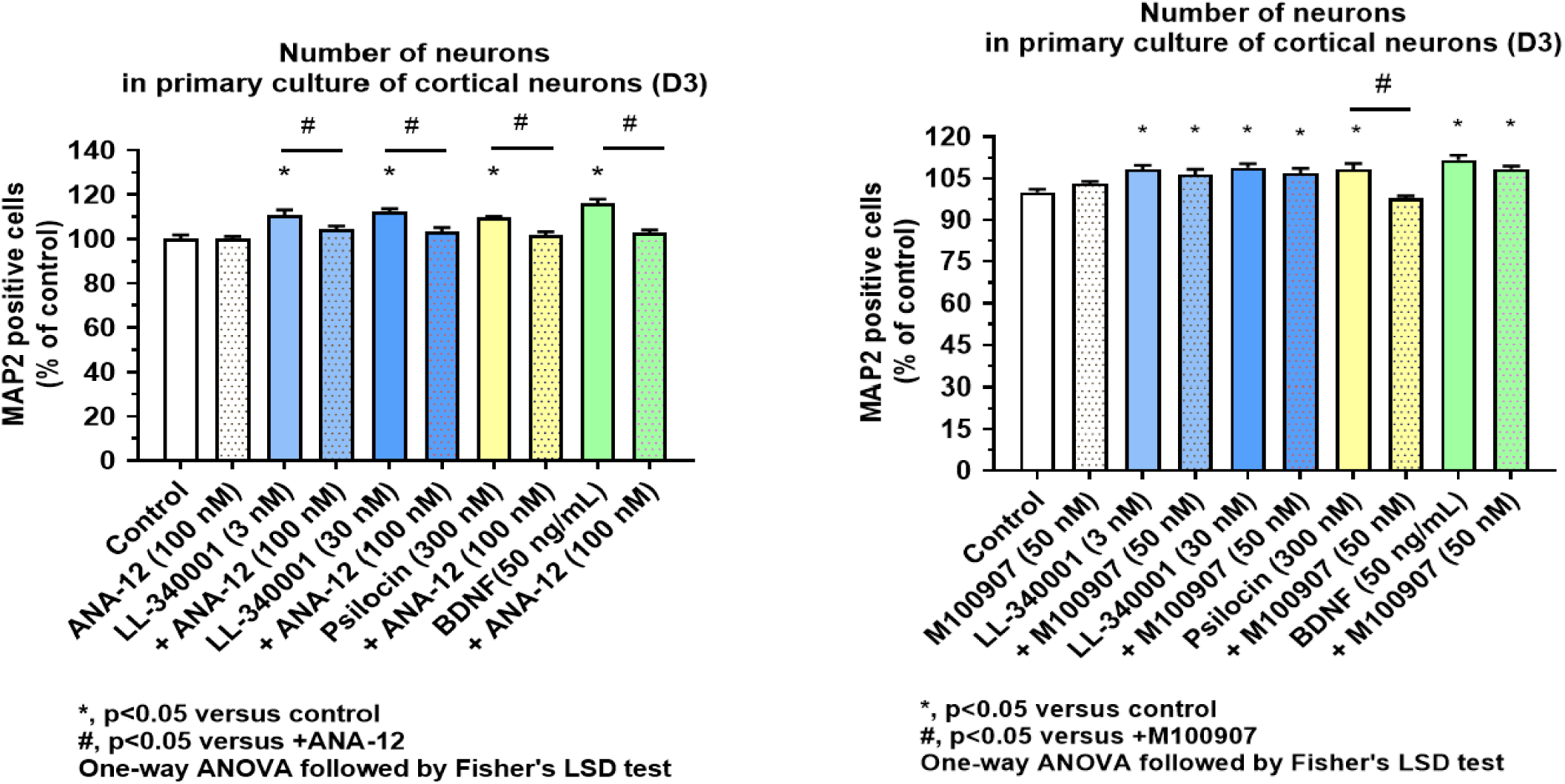
LL-340001-induced cortical neurogenesis requires TrkB signaling but is independent of 5-HT2A signaling. Effect of LL-340001on the number of MAP-2 (+) neurons in a primary culture of cortical neurons (day 3) in the presence or absence of TrkB antagonist ANA-12 (100 nM) (A) or 5-HT2A antagonist M100907 (50 nM) (B). LL-340001 was assessed at 3 nM and 30 nM along with psilocin (300 nM) and BDNF (50 ng/mL) as positive controls. Results are expressed as a percentage of control condition as mean ± SEM (n = 5-6). Data were compared using a one-way ANOVA followed by Fisher’s LSD test. *p< 0.05 versus control was considered significant.

Elunetirom was evaluated in a recombinant human Fatty Acid Amide Hydrolase (hFAAH) hydrolysis assay. Under these conditions, LL-340001 formation increased over time, corresponding to a formation rate of approximately 0.95 nM/min per ng/µL FAAH, and this conversion was completely blocked by the selective FAAH inhibitor PF-3845, confirming FAAH-dependent hydrolysis. In turn, when both compounds were assessed in perinatal mouse brain slices using a composite score of T3-regulated gene expression, elunetirom and LL-340001 were nearly equipotent, with EC50 values of 1.93 and 2.08 nM, respectively. Together, these data indicate that elunetirom is efficiently converted to LL-340001 in FAAH-containing neural tissue, such that the neuronal activity of elunetirom is most parsimoniously explained by rapid in situ generation of the active thyromimetic species.

### LL-340001 increases MAP-2-positive neuron number and neuritogenesis in immature primary cortical neurons

We examined whether LL-340001 could promote early neuronal plasticity phenotypes in immature primary rat cortical neurons. In this young cortical culture system, LL-340001 significantly increased MAP-2-positive neuron number, neurite network, branching roots, and neurite extremities over a low-nanomolar to low-micromolar concentration range, with a bell-shaped concentration-response across endpoints. Thus, in an immature cortical context, LL-340001 increased neuronal cell-number and neurite-elaboration readouts rather than producing a purely neuritic remodeling phenotype.

BDNF behaved as expected, serving as a robust positive control in this assay, increasing neuron number and all neurite-associated structural endpoints. Notably, the magnitude of the LL-340001 effect was comparable to that of BDNF across all cortical readouts. These findings indicate that receptor-active thyromimetic signaling is sufficient to engage a broad developmental plasticity program in immature cortical neurons, encompassing increased MAP-2-positive neuron number together with elaboration of neuronal processes.

### Elunetirom rapidly promotes neuritogenesis and synaptogenesis in mature hippocampal neurons, with persistence through 72 hours

We then asked whether elunetirom could promote plasticity in an established neuronal network, using mature primary rat hippocampal cultures. In contrast to the immature cortical assay, this system was designed to detect remodeling within an established neuronal network rather than changes in neuronal cell number. Elunetirom produced rapid neuritogenic and synaptogenic effects evident within 24 hours and persisting through 72 hours, within an optimal low-nanomolar concentration range.

At 24 hours, 0.3 and 1 nM elunetirom significantly increased neurite network and significantly increased synapse number per neuron at 0.3, 1, and 3 nM. At 72 hours, the neuritogenic effect persisted, with a significant increase in neurite network at 0.3 nM, while synapse number remained elevated at 0.3 and 1 nM. Across both time points, there were no meaningful changes in MAP-2-positive neuron number, consistent with the mature state of the culture and supporting interpretation of the phenotype as neuritic and synaptic remodeling rather than a change in neuronal cell number. BDNF served as a positive control and produced increases in neurite network and synapse number, and the low-nanomolar effects of elunetirom were comparable to those of BDNF.

Importantly, the concentration-response profile in mature hippocampal neurons was bell-shaped, with activity concentrated in the low-nanomolar range. Considered together with the thyroid hormone receptor FAAH prodrug conversion, and brain-slice pharmacology results, these data support the view that elunetirom acts in hippocampal culture through rapid conversion to LL-340001 and subsequent activation of a potent TR-driven neuroplasticity program that is rapidly engaged and sustained over time.

### LL-340001 preserves synapse number under amyloid-associated pathological stress

To determine whether the neuroplastic effects of LL-340001 extended to a pathologically stressed neuronal system, mature hippocampal cultures were pretreated with LL-340001 for 4 hours and then challenged with the amyloid fragment, Aβ1-42 (1.25 µM), and maintained for 24 hours. Under these conditions, LL-340001 produced a significant synaptoprotective effect, preserving synapse number at 1 nM, 3 nM, and 10 nM. The magnitude of this protection was comparable to that produced by BDNF.

The protective effect was confined to synapse number, with no meaningful rescue of MAP-2-positive neuron number or neurite network under the conditions tested. As in the no-insult hippocampal assay, the concentration-response was bell-shaped, with protection observed in an optimal low-nanomolar range and diminished benefit at higher concentrations. These data indicate that receptor-active thyromimetic signaling can preserve synaptic structure under pathological stress, even in the absence of a broader rescue of overall neuritic architecture over this time frame.

### LL-340001 enhances mitochondrial biogenesis-related signaling and mitochondrial output in mature hippocampal neurons

Because sustained neuronal plasticity is tightly coupled to cellular bioenergetics, we next examined whether LL-340001 engaged a mitochondrial biogenesis program in mature hippocampal neurons. After 72 hours of treatment, LL-340001 significantly increased both nuclear PGC-1α and nuclear NRF2 at both 0.3 and 3 nM, indicating activation of transcriptional programs linked to mitochondrial biogenesis and oxidative stress adaptation. At the same concentrations, LL-340001 also significantly increased the number of functional mitochondria, as measured by MitoTracker in MAP-2-positive neurons.

Consistent with these structural and transcriptional changes, LL-340001 increased ATP content, with a numerical increase at 0.3 nM and a significant increase at 3 nM. Together, these results indicate that, in addition to promoting neuritic and synaptic growth, LL-340001 engages an intracellular bioenergetic program that may support these plasticity-related changes.

### LL-340001-induced cortical MAP-2-positive neuron-number effects are TrkB-sensitive and 5-HT2A-independent

To begin to define the downstream signaling pathways associated with the cortical effects of LL-340001, we examined pharmacologic inhibition of TrkB and 5-HT2A signaling in the immature cortical assay. Pretreatment with ANA-12 significantly attenuated the LL-340001-induced increase in MAP-2-positive neuron number at both 3 and 30 nM. In contrast, ANA-12 did not consistently attenuate LL-340001-induced increases in total neurite length, neurite roots, or neurite extremities, indicating that the cell-number and neuritic components of the response were differentially sensitive to TrkB antagonism.

ANA-12 more broadly attenuated the effects of psilocin and BDNF across MAP-2-positive neuron-number and neuritic endpoints.

Pretreatment with the selective 5-HT2A antagonist M100907 blocked the psilocin-induced increase in MAP-2-positive neuron number and reduced its effects on neuritic endpoints, but did not block the effects of LL-340001; BDNF responses were also largely retained.

Neither ANA-12 nor M100907 meaningfully altered baseline cortical neuron number or neurite morphology when administered alone. Taken together, these data support a TrkB-sensitive component of the LL-340001-induced cortical cell-number effect, whereas the neuritogenic effects of LL-340001 were not consistently sensitive to TrkB antagonism and did not require 5-HT2A signaling under the tested conditions.

## Discussion

The present studies identify elunetirom and its active metabolite, LL-340001, as a brain-directed thyromimetic system with neuroplasticity-promoting properties across multiple primary neuronal preparations. The overall mechanism that emerges is coherent: LL-340001 is the receptor-active species, elunetirom is efficiently converted to LL-340001 in FAAH-containing neural tissue, and this active thyromimetic drives downstream remodeling that includes increased MAP-2-positive neuron number and neuritogenesis in immature cortical cultures, rapid neuritic and synaptic remodeling in mature hippocampal neurons, synaptic resilience under pathological stress, and mitochondrial biogenesis-associated signaling. In this framework, elunetirom and LL-340001 share a single functional pharmacology in neuronal systems because elunetirom is rapidly hydrolyzed in situ to LL-340001, the active drug.

This interpretation is biologically consistent with the established role of thyroid hormone signaling in brain development and plasticity [19]. Thyroid hormone is a major regulator of neuronal differentiation, maturation, synaptogenesis, and circuit assembly, and studies in developing and mature neural systems have linked thyroid hormone action to BDNF-dependent structural plasticity. In injured hippocampal slices, thyroxine upregulates BDNF and promotes neuronal survival, whereas in primary hippocampal mouse neurons, thyroid hormone deficiency reduces dendritic arborization, BDNF expression, and TrkB signaling, an effect rescued by exogenous BDNF. More broadly, thyroid hormone receptor signaling is a well-established determinant of neuronal maturation and circuit assembly in the CNS [34–36].

These results extend the mechanistic bridge linking thyroid hormone receptor activation to downstream neuroplasticity through both neurotrophic and bioenergetic effectors, consistent with the central role of BDNF–TrkB signaling in depression and antidepressant-associated plasticity [37, 38]. The inhibitor studies indicate that the LL-340001-induced increase in MAP-2-positive neuron number in immature cortical cultures is sensitive to TrkB inhibition, whereas neurite length, root number, and neurite extremities were not consistently attenuated by ANA-12. Because 10 ng/mL BDNF was present in the standard medium for all vehicle and treatment groups, this basal supplement was controlled across conditions, whereas 50 ng/mL BDNF served as the assay’s optimized positive control for increasing MAP-2-positive neuron number. Accordingly, attenuation by ANA-12 supports reliance on available TrkB signaling for the LL-340001-induced cell-number effect.

Although BDNF abundance was not measured in this experiment, thyroid hormone receptor agonism is known to upregulate BDNF, supporting BDNF-TrkB signaling as part of the proposed pathway. This pattern suggests that the cell-number and neuritic components of the cortical response are not equally dependent on TrkB signaling under these assay conditions. In parallel, LL-340001 increased nuclear PGC-1α, nuclear NRF2, the number of functional mitochondria measured by MitoTracker, and ATP content in mature hippocampal neurons. PGC-1α- and NRF2-linked pathways are relevant to the mitochondrial and oxidative-stress biology increasingly implicated in mood disorders. We therefore interpret these mitochondrial findings not as a parallel observation, but as supportive evidence that LL-340001 engages a coupled neurotrophic and bioenergetic program capable of sustaining neuritic and synaptic remodeling. Importantly, the observed increases in PGC-1α and NRF2, together with concordant increases in the number of functional mitochondria and ATP content, support engagement of a mitochondrial biogenesis- and oxidative stress-response program in mature hippocampal neurons.

Although thyroid-serotonergic crosstalk has been described in other settings, the present data suggest that the neuroplasticity-promoting effects of LL-340001 in these assays do not require 5-HT2A signaling, further supporting a mechanistically distinct, brain-directed thyromimetic pathway. M100907 blocked the psilocin-induced increase in MAP-2-positive neuron number and reduced its neuritic effects but left the effects of LL-340001 intact. Together with the selective sensitivity of the LL-340001-induced cell-number effect to ANA-12, these findings support involvement of TrkB-linked biology without demonstrating direct TrkB engagement by LL-340001 [39–43].

The distinction between the cortical and hippocampal phenotypes is also informative. In immature cortical neurons, LL-340001 increased MAP-2-positive neuron number together with neurite outgrowth, branching roots, and neurite extremities, consistent with a broad developmental plasticity program in an immature culture context. By contrast, in mature hippocampal cultures, the dominant phenotype was rapid neuritic and synaptic remodeling without a meaningful increase in neuron number. These differences are best understood as reflecting neuronal maturity and assay context rather than different primary mechanisms. Early cultures are permissive of changes in neuronal cell-number readouts, whereas established hippocampal cultures are better suited to reveal remodeling of existing neurites and synapses. Readout selection also reflects this difference: branching roots and neurite extremities were measured explicitly in the cortical assay, whereas in the mature hippocampal assay those finer structural changes are effectively captured within the broader neurite-network endpoint.

The bell-shaped concentration-response observed across several assays is likewise consistent with a potent neuroplastic pharmacology operating within an optimal window rather than a simple monotonic dose-response. Similar non-monotonic profiles have been described in neuronal survival and neurite outgrowth systems, particularly for biologically active ligands that engage adaptive cellular pathways [44, 45]. In the present case, the key point is that the most beneficial range repeatedly localized to low nanomolar concentrations, whether the endpoint was increased cortical MAP-2-positive neuron number and neuritic complexity, hippocampal neuritic/synaptic remodeling, synaptoprotection under Aβ challenge, or mitochondrial biogenesis-associated signaling. That repeated concentration window strengthens the view that these are biologically integrated effects of the same active pharmacology.

The effects of elunetirom are especially noteworthy for their speed of onset. Elunetirom increased neurite network and synapse number within 24 hours, with persistence through 72 hours, and these effects were comparable to those of BDNF within an optimal low-nanomolar range. Although a rapid structural response might initially seem surprising for a thyroid hormone receptor-mediated mechanism, it is compatible with the broader pharmacology of thyroid hormone signaling. The brain-slice target-engagement work indicates that T3-regulated genes can be induced within hours, suggesting that receptor engagement can initiate a transcriptional cascade on a timescale compatible with rapid downstream structural remodeling. In this regard, a rapidly acting TR-driven plasticity program may be more plausible than traditionally assumed.

From a translational standpoint, the combination of rapid synaptogenic effects, mitochondrial remodeling, and TrkB-linked plasticity is consistent with a potentially rapid-acting antidepressant mechanism. Convergent evidence in major depressive disorder implicates impaired hippocampal and cortical neuroplasticity, reduced synaptic integrity, and dysregulated BDNF-TrkB signaling, while rapid-acting antidepressant interventions are increasingly understood through the lens of structural and synaptic plasticity [46, 47].

Bipolar disorder also shows strong links to altered BDNF biology and mitochondrial dysfunction, making a mechanism that couples neurotrophic and bioenergetic remodeling potentially relevant to bipolar depression as well as MDD [48–50]. Against that backdrop, the present in vitro profile suggests that brain-directed thyromimetic signaling may offer a differentiated route to antidepressant-relevant plasticity. A useful clinical comparator for a more scalable, rapidly acting, mechanistically differentiated antidepressant strategy is oral dextromethorphan-bupropion, which is approved for MDD, whereas intranasal esketamine remains tied to supervised administration under a REMS framework; mechanistic discussions place dextromethorphan-containing antidepressant strategies within the broader non-monoaminergic, plasticity-relevant space through NMDA receptor modulation, sigma-1 receptor signaling, and downstream network biology relevant to trophic and plasticity responses [22–24].

Finally, the Aβ study extends the mechanism from basal plasticity to synaptic resilience under pathological stress. In this setting, LL-340001 preserved synapse number without materially altering overall neuron number or neurite network over the same interval, suggesting that synapses may be an especially sensitive early target of thyromimetic protection. That finding strengthens elunetirom’s translational relevance by showing that the same pharmacology that enhances basal synaptic remodeling can also oppose synaptic loss in a stressed neuronal system. Future work will explore whether this resilience generalizes to other pathophysiologically relevant insults, including glutamatergic and glucocorticoid stress paradigms.

## Acknowledgments

The authors thank Neuro-Sys SAS (Gardanne, France) for conducting the primary cortical and mature hippocampal neuronal culture studies, including culture preparation, compound treatment, immunocytochemical staining, image acquisition and analysis, and provision of the associated study data and reports. The authors also acknowledge the Neuro-Sys scientific and technical teams for their contributions to study execution and technical discussions.

